# Academic Success and Mental Health: The Paradox of Frontoparietal-Default Mode Network Coupling among Children Facing Poverty

**DOI:** 10.1101/2024.10.18.619123

**Authors:** Selina Pacheco, Silvia A. Bunge, Monica E. Ellwood-Lowe

**Author notes:** Corresponding Author Monica E. Ellwood-Lowe, Department of Psychology University of Pennsylvania, United States.

## Abstract

Childhood family income is a powerful predictor of academic achievement and mental health. Here, we ask whether children living in poverty who succeed academically are subsequently protected from, or at risk for, internalizing symptoms. Prior research indicates that children in poverty with better academic performance tend to have higher temporal coupling between the Lateral Frontoparietal Network (LFPN) and Default Mode Network (DMN) than lower-performing children in poverty. An open question is whether higher LFPN-DMN coupling has maladaptive long-term consequences for mental health for this population. In this pre-registered longitudinal study, we analyzed data from 10,829 children (1,931 in poverty) in the ABCD study across four time points (ages 9-13). Higher grades correlated with fewer internalizing symptoms; this association was more pronounced for children below poverty. Longitudinally, LFPN-DMN connectivity correlated positively with internalizing symptoms across both groups and timepoints. Thus, although higher academic performance was associated with better mental health outcomes for all children, the specific pattern of LFPN-DMN connectivity that supports academic resilience among children in poverty may be a risk factor for developing internalizing symptoms. These findings highlight the complex nature of academic resilience in the context of structural inequity.

**Highlights:**

- High grades linked to fewer internalizing symptoms, especially for kids in poverty.
- High LFPN-DMN connectivity predicts higher internalizing symptoms in children.
- Neural correlates of academic resilience may predispose children to internalizing.
- Children below poverty had higher internalizing symptoms.

## Introduction

Childhood family income is a powerful predictor of academic achievement and mental well-being. For millions of children born into poverty, the path to academic success is laden with obstacles, yet some manage to defy the odds and excel academically. Previous research suggests that academic success offers hope to children from impoverished backgrounds by providing them with a pathway out of poverty (UNESCO, 2017). But does succeeding academically come at a cost? Here, we ask whether children living in poverty who defy the odds by succeeding academically are subsequently protected from–or alternatively more vulnerable to–internalizing disorders.

In the United States alone, approximately 11.3 million children and adolescents grapple with the harsh realities of poverty, facing limited access to resources and opportunities (Shrider et al., 2021). These socioeconomic disadvantages not only hinder educational prospects but also exacerbate mental health issues, often leading to higher rates of anxiety, depression, and behavioral problems (Reiss, 2013). Despite the systemic barriers they encounter, some children in poverty achieve academic success comparable to their more affluent peers. Intriguingly, previous research has shown that these academically resilient children from low-income backgrounds have different brain connectivity patterns compared to those typically linked with high cognitive performance. This challenges the conventional belief that poverty is always associated with cognitive deficits (Ellwood-Lowe et al., 2021; Merz et al., 2024). Here, we follow up on this finding to ask: what are the implications of these neural differences for the mental health outcomes of academically resilient children living in poverty?

Risk and resilience models in child development provide a framework for understanding these complex dynamics, highlighting how some children manage to thrive despite facing significant adversities (McDorman et al., 2024). These models emphasize the interplay between individual traits and environmental factors, underscoring the importance of protective factors that can mitigate the negative impacts of poverty.

## Impact of Socioeconomic Status on Academic Achievement and Mental Health

### Academic Achievement

Socioeconomic status (SES) plays a critical role in shaping children’s academic outcomes, influenced both by individual and structural factors. Children from low-SES families often attend underfunded schools with fewer resources, less experienced teachers, and larger class sizes. These schools may lack advanced coursework, extracurricular activities, and adequate facilities, creating an environment that hinders academic achievement (Pfeffer, 2018). Neighborhood factors, such as high crime rates and lack of safe recreational spaces, further exacerbate these challenges by limiting children’s opportunities for cognitive and social development (Aikens & Barbarin, 2008).

Additionally, low-SES families may face housing instability, food insecurity, and limited access to healthcare, all of which negatively impact children’s cognitive functioning and school performance (Gershoff et al., 2007). Chronic stress from financial instability and adverse living conditions can impair attention, memory, and executive function, essential for learning and academic success (Harms & Garrett-Ruffin, 2023). Moreover, early language exposure significantly influences later linguistic skills and cognitive abilities, with greater conversational turns with adults linked to enhanced activation in Broca’s area, critical for language processing, highlighting the importance of environmental factors on cognitive development (Romeo et al., 2018; Merz et al., 2020; Lurie et al., 2021). These educational challenges underscore the significant impact of socioeconomic status on academic achievement beyond individual characteristics of caregivers. However, the consequences of low SES extend beyond the classroom, profoundly influencing children’s mental health as well.

### Mental Health

Economic hardship can lead to chronic stress, which can negatively impact brain development and emotional regulation in children (Coley et al., 2015). This chronic stress not only impairs cognitive functions such as attention, memory, and executive function, essential for emotional well-being and mental health, but also contributes to lasting deficits in mental health issues such as depression, anxiety, and behavioral problems throughout life (McLeod & Shanahan, 1996; Reiss, 2013; Rosen et al, 2019). Additionally, socioeconomic disadvantage can affect brain structure and function, including reduced gray matter and altered activation in areas related to language and executive function, which contribute to mental health outcomes (Merz et al., 2024). Children in low-SES families are more likely to experience adverse childhood experiences (ACEs) such as neglect, abuse, and exposure to violence, all of which are significant risk factors for developing mental health disorders. These adverse experiences can have a cumulative effect, exacerbating mental health issues as children grow older (Walsh et al., 2019). Recent research has highlighted that different types of childhood adversity, specifically deprivation and threat, have distinct impacts on cognitive and emotional development even in early childhood. Deprivation, associated with a lack of cognitive and social inputs, is linked to worse cognitive control, while threat, linked to harm or the threat of harm, is associated with altered fear learning and blunted physiological reactivity (Machlin et al., 2019).Notably, parent emotion regulation plays a differential role in how children cope with early adversity, impacting their own emotion regulation abilities (Milojevich et al., 2021).

### Interconnected Effects

It is crucial to acknowledge that children growing up in low-income households are a diverse group, and while some individuals manage to excel academically despite facing considerable obstacles, the mainstream narrative often emphasizes individual achievement without adequately addressing systemic barriers. These barriers, such as limited access to quality education, economic instability, and lack of support systems, create immense challenges for many children from low-income backgrounds. This raises an important question: does academic success under such challenging circumstances ultimately come at a cost to an individual’s mental health?

One study offers intriguing insights into this question. The authors followed individuals with exceptionally successful careers, and found that these people were not only medically but also psychologically better off, pointing to a positive link between achievement and mental health (Kell et al., 2022). However, a closer examination of their methodology reveals potential limitations. Participants were primarily drawn from a sample of adults identified as having exceptionally high test scores at a young age, and success was defined based on their adult income. Notably, this recruitment strategy may have skewed the sample towards individuals who did not encounter significant structural barriers to success. Moreover, the demographic composition of the participants, predominantly white and likely growing up in relatively affluent environments, underscores the need to consider intersectional factors such as race and socioeconomic background in understanding the relationship between success and mental health.

Other studies have explored how resilience might mitigate the negative effects of childhood poverty on health. For instance, one study explored the role of resilience in mitigating the negative effects of childhood poverty on health (De France et al., 2022). Their research assessed physiological stress and ill health using a composite index termed “allostatic load,” which captured the cumulative impact of stress on the body over time. While they observed that low-income children with high self-control, self-esteem, and engaged parenting demonstrated normal psychological adjustment, they also found a concerning trend. These resilient children, while not displaying evident mental health problems, appeared to pay a physical toll for their adaptive coping mechanisms. They experienced greater physiological stress and ill health, which could over time result in a higher disease burden, such as an increased incidence of cardiovascular disease (McEwen, 1998; Cohen et. al, 2007; Chrousos, 2009).

The relationship between socioeconomic status (SES), academic achievement, and mental health lays the groundwork for understanding the complex interplay of structural factors and individual outcomes. We see that while some individuals may thrive academically and enjoy favorable psychological outcomes, others may endure hidden costs, trading off psychological resilience for physical well-being. However, less work has examined neural mechanisms underlying these associations, especially from a developmental perspective. By exploring how academic resilience manifests at the neural level in childhood, we can gain insight into the cognitive and socioemotional processes that shape academic success and mental well-being among individuals from different socioeconomic backgrounds.

## Brain Networks in Relation to Socioeconomic Status, Academic Achievement and Mental Health

### Default Mode Network, Lateral Frontoparietal Network, and Cingulo-Opercular Network

We used Functional Magnetic Resonance Imaging (fMRI) when participants were at rest to investigate the brain connectivity patterns associated with academic resilience in children from low-income backgrounds. This non-invasive technique that measures and maps brain activity by detecting changes in blood flow (Ogawa et al., 1990; Van den Heuvel & Pol, 2010). When a brain region is more active, it consumes more oxygen, and to meet this increased demand, blood flow to that region increases. fMRI captures these changes, providing insights into the brain’s functional architecture and connectivity. Here, we focus on networks of brain regions that typically coactivate together, defined from resting state fMRI (Van den Heuvel & Pol, 2010; Gordon et al., 2017). Understanding these patterns can help elucidate the cognitive and socioemotional processes that underlie academic success and mental well-being.

In this study, we focus on three specific brain networks: the default mode network (DMN), the lateral frontoparietal network (LFPN), and the cingulo-opercular network (CON). These networks serve different cognitive functions and have been implicated in both cognitive and socioemotional success for children. The DMN is activated when the brain is at rest or not focused on something outside of the here-and-now, for example during introspection, self-referential thinking, and mind wandering (Christoff et al., 2016; Spreng et al., 2010; Raichle et al., 2001). In contrast, the LFPN is involved in executive functions, including attention, working memory, reasoning, and decision-making. It plays a significant role in top-down attentional and cognitive control, which allows individuals to concentrate on relevant information while disregarding irrelevant stimuli (Seeley et al., 2007; Vincent et al., 2008; Niendam et al., 2012). The LFPN and the DMN are generally thought to play contrasting roles in cognition, in a dynamic interplay between top-down attentional control and bottom-up sensory processing (Corbetta & Shulman, 2002; Fox et al., 2005; Andrews-Hanna et al., 2014). Additionally, the CON has been theorized to act as a mediator between the DMN and the LFPN, signaling the LFPN when a controlled response is needed and facilitating the transition from the DMN’s “default” mode to a state of top-down control (Dosenbach et al., 2007; Sridharan et al., 2008; Uddin et al., 2011).

Overall, there is evidence that lower connectivity between the LFPN and the DMN is related to better cognitive performance and mental health (Chai et al., 2014; DeSerisy et al., 2021; Lopez et al., 2020; Sherman et al., 2014; Whitfield-Gabrieli et al., 2020; Belleau et al., 2023). Longitudinal research indicates that higher connectivity between the LFPN and DMN in childhood is associated with worse mental health outcomes (Chai et al., 2014; DeSerisy et al., 2021; Lopez et al., 2020; Shapero et al., 2019; Sherman et al., 2014; Whitfield-Gabrieli et al., 2020; Belleau et al., 2023). However, these studies were largely conducted with small convenience samples, leaving the generalizability to individuals in poverty unclear.

In contrast to the research with generally higher-SES children, we previously reported different associations between LFPN-DMN connectivity and cognitive and academic performance for children above and below poverty in the Adolescent Brain Cognitive Development (ABCD) study (Ellwood-Lowe et al., 2021; 2022). Specifically, children in poverty with better cognitive test performance and grades tended to have *higher* coupling between LFPN and DMN than lower-performing children in poverty, in direct contrast to the pattern observed for children above poverty. Thus, an open question is whether this pattern of connectivity, which has been linked to higher achievement for children in poverty, might have maladaptive long-term consequences, particularly for mental health.

### The Present Study

Building upon previous research, our study investigates how high academic achievement and associated brain function may influence mental health outcomes, specifically focusing on internalizing symptoms in children from low-income backgrounds. To address these gaps we analyzed data from 10,829 children (1,931 in poverty) in the ABCD study at baseline (ages 9-10 years) and followed them through four time points over three years. Our study aims to examine the predictive relationships between academic performance, LFPN-DMN connectivity, CON-DMN connectivity, and CON-LFPN connectivity strength, and internalizing symptoms. Furthermore, we ask whether these associations differed between children above and below the poverty line, contributing to a nuanced understanding of the role of socioeconomic status in shaping neural correlates of academic achievement and mental health outcomes.

## Methods

Analysis plans were pre-registered prior to analyzing the data (https://aspredicted.org/CKB_C11) and analysis scripts will be available on the Open Science Framework. The original data are available with permissions on the NIMH Data Archive (https://nda.nih.gov/abcd).

### Participants

Participants were part of the ABCD study (Barch et al., 2018; Jernigan et al., 2018). To ensure the study’s findings are relevant to the entire U.S. population, participants were selected to reflect diverse demographics including various races, ethnicities, education levels, income levels, and living environments. Recruitment efforts involved a partnership with public, private, and charter elementary schools across the continental United States. Researchers identified schools within a reasonable distance from each of the 21 research sites and selected a subset for participation. This selection process was staggered over two years to manage outreach effectively. Principals and administrators of these schools distributed information packets to families of 8- to 10-year-old students, using school folders, postal mail, or email lists provided by the study at no cost to the schools. Informed consent was obtained from all participating families. Additionally, the study received Institutional Review Board approval (Barch et al., 2018; Jernigan et al., 2018). While the final sample does not perfectly mirror the national population, these efforts have resulted in one of the most demographically representative neuroimaging studies to date.

We estimated children’s poverty status using the supplemental poverty measure, based on their family income, household size, and the average supplemental poverty level for the study sites included in the sample, since study site data were de-identified. Families with four members or less were classified as living in poverty if their total income was below $25,000, while families with five or more members were classified as such if their total income was below $35,000, as further explained in Ellwood-Lowe et al. (2021). We excluded children who did not provide usable data for grades, resting state, and internalizing symptoms. This left us with a total of 8,898 children, of whom 1,931 were identified as likely living below the poverty line at the baseline assessment (see Table 2 for numbers across all timepoints).

**Table 1.**
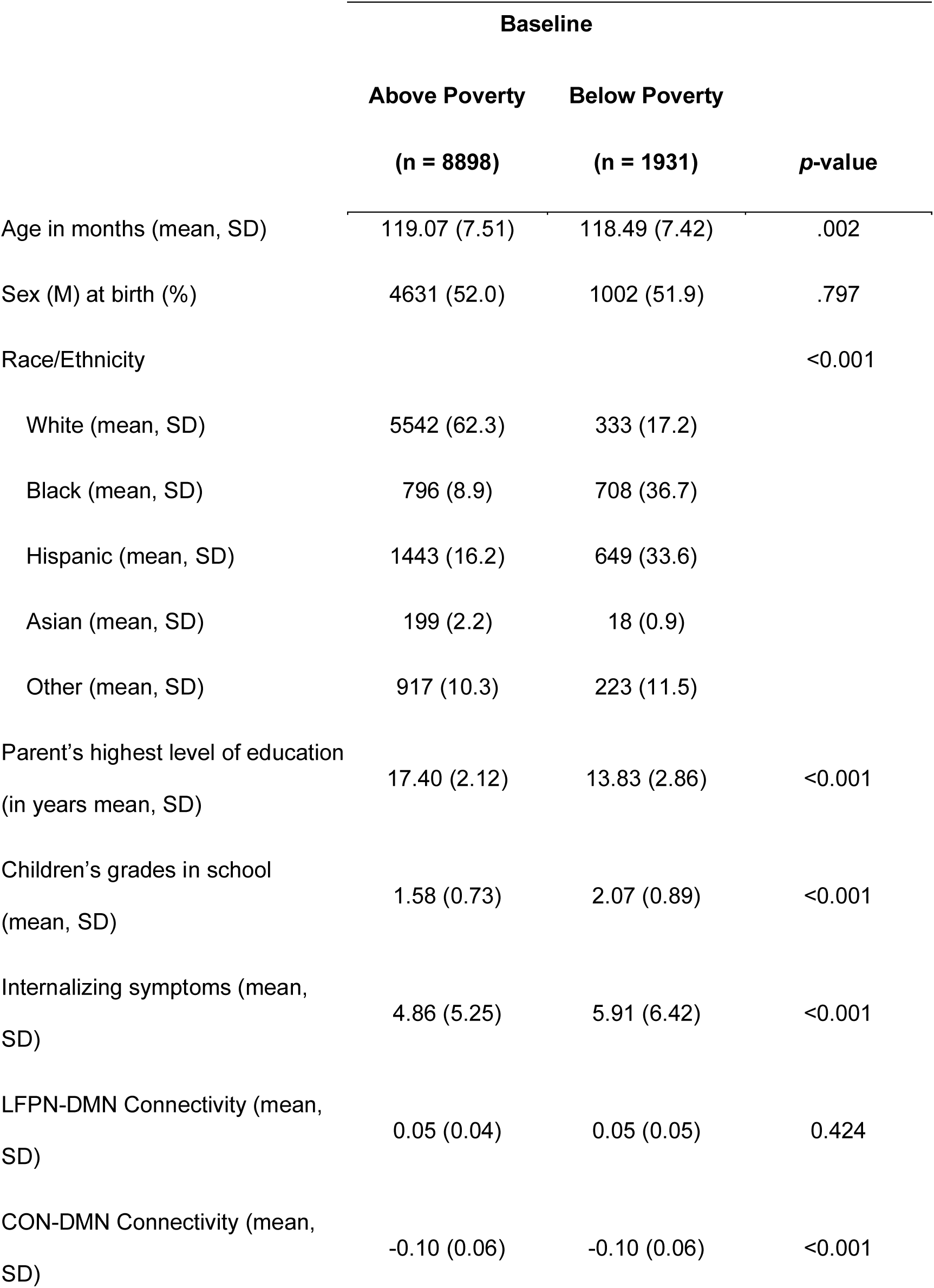
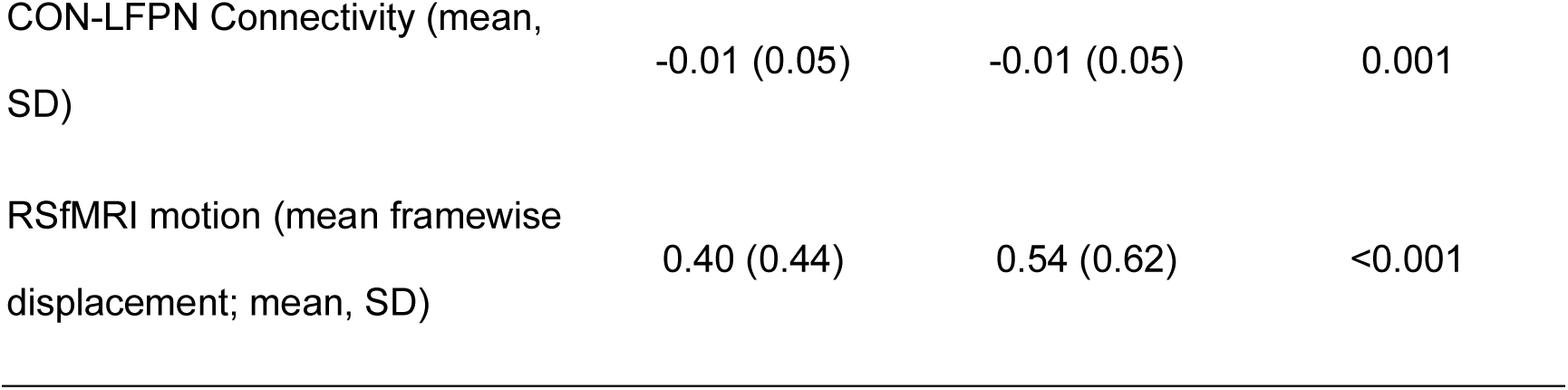
Participant characteristics for children above and below poverty in our final sample. *p*-value based on t-test of difference between groups.

**Table 2.** Usable datapoints across each timepoint in the current study.

| Timepoint |  | Above<br>Poverty | Below<br>Poverty |
| --- | --- | --- | --- |
| **One-year follow-up**<br>**(T1)** | *N* | 8554 | 1716 |
|  | Age | 10.93 (7.74) | 10.88 (7.59) |
| **Two-year follow-up**<br>**(T2)** | *N* | 6312 | 1136 |
|  | Age | 11.97 (7.82) | 11.87 (7.73) |
| **Three-year follow-up**<br>**(T3)** | *N* | 6090 | 1009 |
|  | Age | 12.92 (7.73) | 12.83 (7.74) |

### Grades

Actual school records across the many school districts and sites involved in the ABCD Study were not available at the outset of the study, thus our study relied on youth and caregiver self-report of children’s average grades in school. The responses for academic performance are assigned a value of “1” to the highest possible performance (i.e., A) and the value assigned increases with poorer grades (i.e., B = 2, and so forth, with F = 5).

### Internalizing Symptoms Measures

Children’s internalizing symptoms were measured using the Child Behavior Checklist (CBCL), a part of the Achenbach System of Empirically Based Assessment (ASEB A) used to detect behavioral and emotional problems in children and adolescents (Achenbach, 1999). The CBCL is completed by the caretaker who spends the most time with the child. This produces measures of internalizing symptoms as a whole, as well as symptoms across three subscales: withdrawal/depression, anxiety/depression, and somatic symptoms. The range of CBCL scores in our sample at baseline was 33 - 93 (see Table 1). 722 (6.67%) participants exceeded a clinical threshold of 69 while an additional 410 (3.79%) participants were within the borderline range of 65 - 69 (Achenbach, T.M. & Rescorla, L.A., 2001).

### MRI Scan Procedure

Scans were typically completed on the same day as the cognitive battery, but could also be completed at a second testing session. After completing motion compliance training in a simulated scanning environment, participants first completed a structural T1-weighted scan. Next, they completed 3-4 five-minute resting-state scans, in which they were instructed to lay with their eyes open while viewing a fixation cross on the screen. The first two resting state scans were completed immediately following the T1-weighted scan; children then completed two other structural scans, followed by 1-2 more resting state scans, depending on the protocol at each specific study site. All scans were collected on one of three 3T scanners with an adult-size head coil (Casey et al., 2018). Structural and functional images underwent automated quality control procedures (including detecting excessive movement and poor signal-to-noise ratios) and visual inspection and rating (for structural scans) of images for artifacts or other irregularities (Hagler et al., 2019); participants were excluded if they did not meet quality control criteria, including having fewer than 12.5 min of data with low head motion (framewise displacement < 0.2 mm).

### Scan Parameters and Resting-State fMRI Processing

T1-weighted structural MRI and fMRI scans were acquired from 19 study sites. The T1-weighted scans were collected in the axial position, with 1mm^3^ voxel resolution, 256 × 256 matrix, 8 degree flip angle, and 2x parallel imaging. Although some scan parameters differed between scanner platforms (Siemens, Philips, and GE), they were optimized to be compatible to ensure maximal comparability across the study sites.

The fMRI scans were also acquired in the axial position, with 2.4mm^3^ voxel resolution, 60 slices, 90 × 90 matrix, 216 × 216 field of view (FOV), 800 ms repetition time (TR), 30 ms echo time (TE), 52 degree flip angle, and 6-factor MultiBand Acceleration. Real-time procedures were used to monitor head motion during the scan acquisition and to adjust scanning procedures as necessary. This prospective motion correction procedure was found to significantly reduce scan artifacts due to head motion (Casey et al., 2018).

Preprocessing was carried out by the ABCD Data Analysis and Informatics Core (Hagler et al., 2019). Children’s resting-state fMRI brain networks were identified using the Gordon parcellations, a specific set of brain regions that have been identified as forming function networks based on their connectivity patterns (Gordon et al., 2016). This produced standardized values for the connectivity within and between large-scale networks; here, building on our prior work (REFS), we focus specifically on connectivity between the LFPN, DMN, and CON.

### Analysis

Analyses were performed using R [version 4.2.1]. We were interested in the interaction between poverty status and internalizing symptoms in predicting grades and between-network connectivity. Specifically, we performed 4 separate linear mixed-effects models, as pre-registered, using the lme4 package (Bates et al., 2014) to separately test the relation between (1) grades, (2) LFPN-DMN, (3) CON-DMN, and (4) CON-LFPN connectivity and children’s internalizing symptoms. All models included an interaction of poverty level, fixed effects for child age, and random intercepts for site and family; LFPN-DMN, CON-DMN, and CON-LFPN models additionally included fixed effects for head motion (mean framewise displacement). For each of our major analyses, we pre-registered several different possible patterns of results, as specified below. We focus on analyses with LFPN-DMN connectivity in the main text, and report those with CON-DMN and CON-LFPN connectivity in the Supplement.

#### Association between grades and internalizing symptoms

We hypothesized that higher grades would be associated with fewer internalizing symptoms for children above and below poverty (main effect of grades on internalizing). Additionally, we considered the possibility that the association between grades and internalizing symptoms might differ across groups, such that children in poverty would show a weaker, or even opposite association, such that those students with higher grades would have more internalizing symptoms (interaction between grades and poverty status on internalizing).

#### Associations between connectivity and internalizing symptoms

We hypothesized that higher LFPN-DMN, CON-DMN, and CON-LFPN connectivity would be associated with more internalizing symptoms for children both above and below poverty (main effect of connectivity on internalizing). Additionally, we considered the possibility that the association between one or more of these connectivities and internalizing symptoms might differ for children in poverty. Specifically, we predicted that children in poverty might show a less strong, or even opposite association, such that higher LFPN-DMN, CON-DMN, and/or CON-LFPN connectivity is associated with fewer internalizing symptoms (interaction between connectivity and poverty status on internalizing).

### Longitudinal analyses

Our primary pre-registered analyses focused on these relations at baseline, when children were ages 9-10. While we pre-registered that we would explore longitudinal associations, we did not specify a concrete analysis plan. We adopted two different approaches for longitudinal analyses. First, we examined whether baseline grades and connectivity, respectively, relate to children’s *later* development of internalizing symptoms. To do this, we used linear mixed effects models to assess the relationship between baseline grades and connectivity measures with internalizing symptoms at follow-up, while controlling for internalizing symptoms at baseline and relevant covariates including age, sex, and socioeconomic status; these analyses are reported in the main text. Our second set of analyses tested the association between grades and resting state metrics across all timepoints, while controlling for relevant covariates. We examined longitudinal associations by running the same mixed-effects models as in our first set of analyses, additionally including a time point interaction with grades or connectivity (as appropriate), across four time points, and a random intercept for participants to account for repeated measures. These analyses are included in the supplement for completeness.

### Internalizing subscales

As exploratory analyses, we also examined whether associations for our main research questions varied across the three internalizing subscales (anxiety/depression, withdrawal/depression, and somatic symptoms). These did not differ meaningfully from our main results and are reported in the Supplement for completeness.

## Results

### Association Between Children’s School Performance and Internalizing Symptoms

Across the group, worse grades were associated with greater internalizing symptoms (*B* = 2.00, SE = 0.78; *X*^2^ (4) = 195.55, *p* < .001). Children in poverty experienced greater internalizing symptoms on average than children above poverty (*B* = 1.55, SE = 0.38; *X*^2^ (1) = 8.79 *p* = .003). In addition, the association between internalizing symptoms and grades varied as a function of whether or not children were in poverty (*X*^2^ (4) = 14.18, *p* < .007). Specifically, while children both above and below poverty with worse grades typically had higher internalizing symptoms, this association was stronger for children below poverty (*B* = 5.94, SE = 0.87; *X*^2^ (4) = 64.72, *p* < .001) than for children above poverty (*B* = 2.05, SE = 0.75; *X*^2^ (4) = 137.29, *p* < .001), suggesting a tighter link between grades and internalizing for children in poverty.

We also explored differences as a function of internalizing subscales, as detailed in the supplemental material. For both anxiety/depression and withdrawal/depression, worse grades were linked to higher symptoms, with a stronger effect for children in poverty. For somatic symptoms, worse grades were associated with higher symptoms for all, with no significant interaction based on poverty.

To examine whether children’s grades in school were related to *later* internalizing symptoms, we performed an exploratory analysis of the association between baseline school grades with internalizing symptoms one year later, when controlling for baseline internalizing symptoms. The results indicated that grades were linked to internalizing symptoms cross-sectionally but not longitudinally, when controlling for baseline internalizing (see Supplement).

### Association Between Children’s LFPN-DMN Connectivity Levels and Internalizing Symptoms

Next, we examined the relation between children’s LFPN-DMN connectivity and their internalizing symptoms at baseline (Figure 2). We found a main effect of LFPN-DMN connectivity in predicting children’s internalizing symptoms (*B* = 4.61, SE = 1.45; *X*^2^ (1) = 9.88, *p* = .002), indicating that children who had higher LFPN-DMN connectivity typically had higher internalizing symptoms. However, the interaction between LFPN-DMN connectivity and children’s poverty status did not reach significance (*X*^2^ (1) = 0.57, *p* = .451), suggesting the association did not differ significantly as a function of children’s poverty status. Thus, higher LFPN-DMN connectivity, while potentially cognitively adaptive for children in poverty, may be a risk factor for the development of internalizing symptoms for all children.

**Fig. 1.**
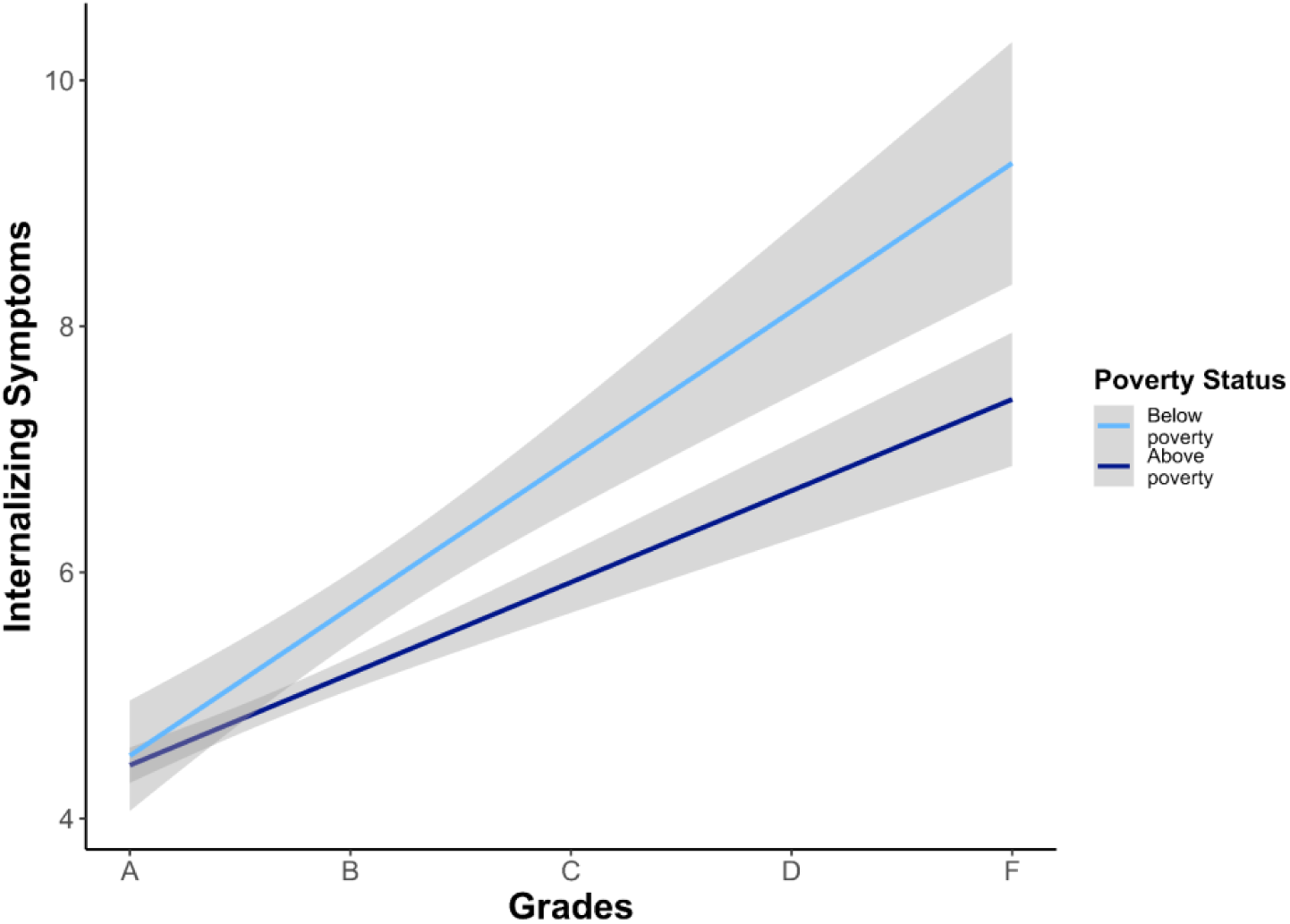
Association Between Children’s School Grades and Their Internalizing Symptoms. Relation between children’s school grades and their internalizing symptoms for children below poverty (light blue) and children above poverty (dark blue). The gray bars indicate mean values with ±95% confidence intervals for a linear model.

**Fig. 2.**
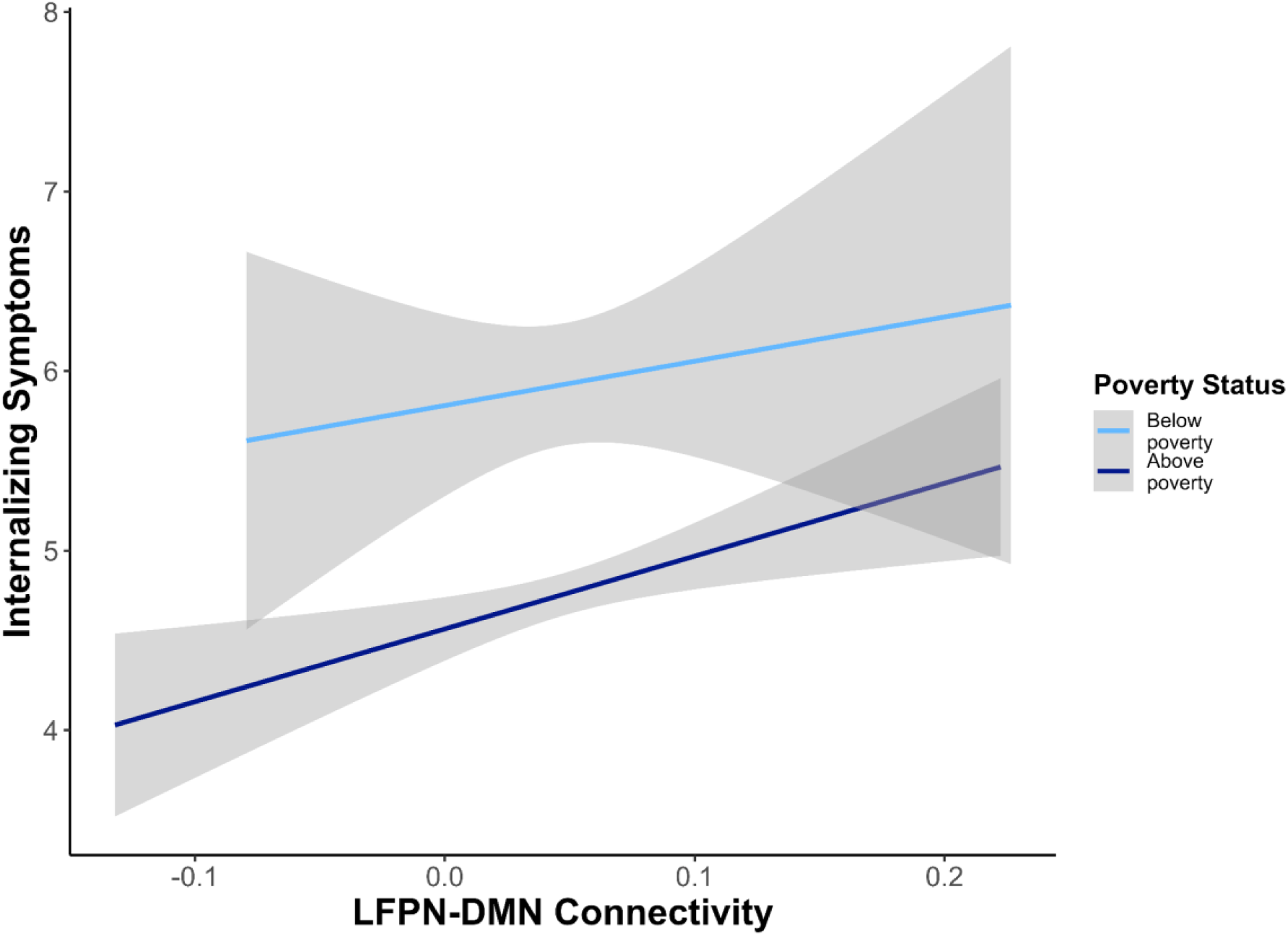
Association Between Children’s LFPN-DMN Connectivity Levels and Internalizing Symptoms. Relation between children’s LFPN-DMN connectivity levels and their internalizing symptoms for children below poverty (light blue) and children above poverty (dark blue). The gray bars indicate mean values with ±95% confidence intervals for a linear model.

We also explored differences by internalizing subtypes, as detailed in the supplemental material. For anxiety/depression and withdrawal/depression symptoms, higher LFPN-DMN connectivity was linked to higher symptom severity, regardless of poverty status. For somatic symptoms, there was no significant main effect or interaction.

To examine whether this pattern of brain connectivity was related to *later* internalizing symptoms, we next conducted a longitudinal analysis examining the association between baseline LFPN-DMN connectivity with internalizing symptoms one year later, when controlling for baseline internalizing symptoms and poverty status. Although the preregistration did not specify the precise approach for the longitudinal analyses, this question was central to our investigation. While there was no consistent association between LFPN-DMN and later internalizing symptoms across the group, there was a significant interaction between LFPN-DMN connectivity and poverty status (Table 3). Breaking down this interaction revealed that higher connectivity was associated with more internalizing symptoms one year later for children in poverty in particular (*B* = 5.83, SE = 2.76, *X^2^* = 4.46, *p* = .035), but not for children above poverty (*B* = -0.90, SE = 1.07, *X^2^* = 0.70, *p* = .402). This finding highlights that the relationship between LFPN-DMN connectivity and later internalizing symptoms may be contingent upon children’s socioeconomic context, given that higher LFPN-DMN connectivity predicts future internalizing symptoms only for children below the poverty line in this large sample.

**Table 3.**
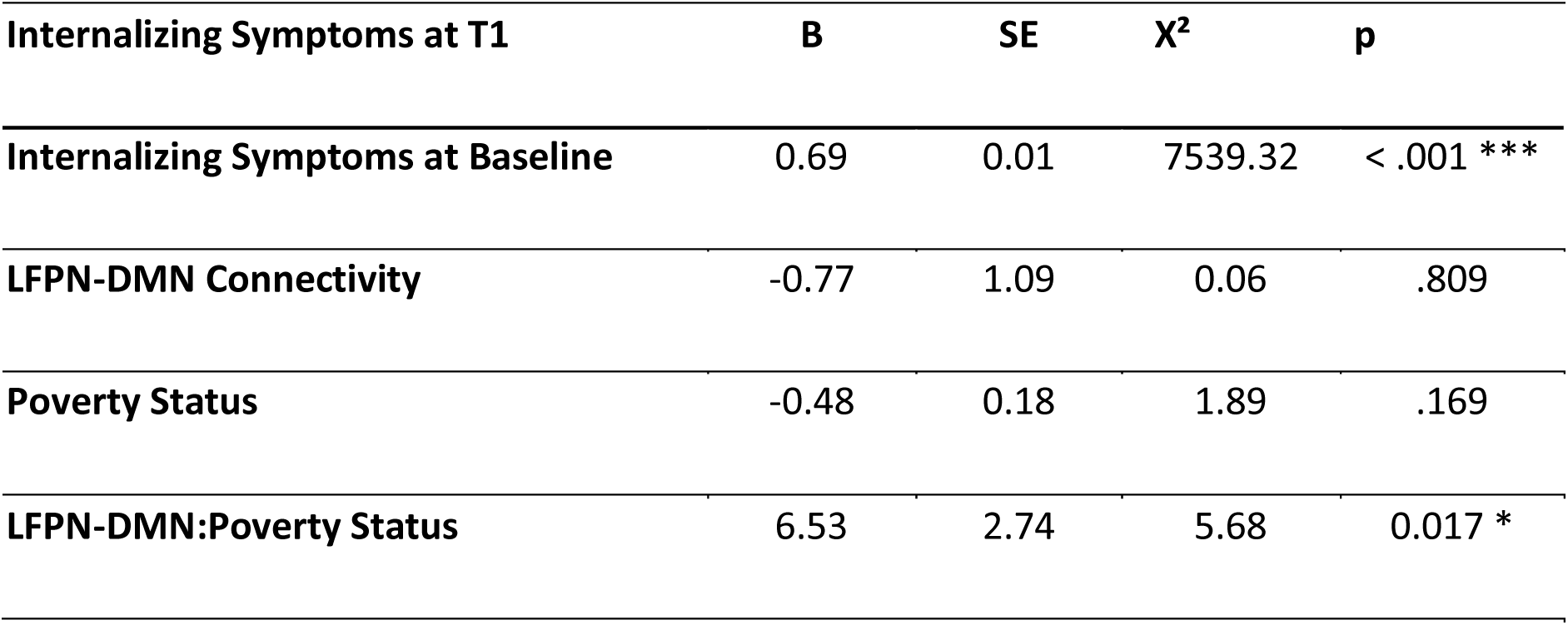
Results of linear mixed effects model associating internalizing symptoms at T1 with a two-way interaction between LFPN-DMN connectivity and poverty status at T1 while controlling for internalizing symptoms at baseline. Chi-squared and significance values from Type II anova, using the Anova function in *car*.

### Association Between Children’s CON Connectivity and Internalizing Symptoms

We next examined the relation between CON-DMN and CON-LFPN connectivity and internalizing symptoms at baseline. None of the cross-sectional or longitudinal analyses were significant. Children’s CON-DMN connectivity levels and CON-LFPN connectivity levels did not show a significant association with internalizing symptoms, nor was there a significant interaction with poverty status.

## Discussion

In the present study, we aimed to investigate relationships between children’s school performance, brain connectivity, and internalizing symptoms, with a focus on the role of poverty. Previous research has shown that high-achieving children from lower-income backgrounds exhibit different patterns of brain connectivity than their higher-income counterparts (Ellwood-Lowe et al., 2021). Here, we examined the relationship between children’s grades in school and their internalizing symptoms, and LFPN-DMN, CON-DMN, and CON-LFPN connectivity, and whether these differed for children in poverty. This allowed us to investigate whether high academic achievement in children living in poverty, along with its associated brain connectivity patterns, might be risk factors for poor mental health.

We found that worse grades were associated with greater internalizing symptoms, and this association was stronger for children living below the poverty line. Additionally, higher LFPN-DMN connectivity (but not connectivity between pairs of networks involving the CON) was linked to higher internalizing symptoms across all children, with longitudinal analysis revealing that this connectivity predicted future internalizing symptoms specifically for children in poverty. While grades were negatively correlated with internalizing symptoms, they did not predict future internalizing symptoms; by contrast, functional brain connectivity did. This suggests that while academic performance and mental health are linked at any given time, grades alone are not reliable indicators of future emotional well-being. One possible interpretation is that grades reflect the current state of how children are managing their academic and emotional challenges but do not capture the neurodevelopmental mechanisms that drive longer-term mental health outcomes.

Our findings align with previous research indicating that academic performance is closely tied to emotional well-being in children. For instance, prior studies have shown that lower academic achievement is often associated with higher levels of anxiety and depression (Duncan et al., 2012; Bas, G., 2021; Moilanen et al., 2010). However, our results extend this understanding by highlighting the exacerbated impact on children in poverty, suggesting that socioeconomic factors may intensify the relationship between academic struggles and internalizing symptoms. This is consistent with theories of cumulative risk, which propose that children facing multiple adversities are more likely to experience negative outcomes (Hodgkinson et al., 2017; Evans et al., 2013; Dashiff et al., 2009).

Regarding brain connectivity, our findings that higher LFPN-DMN connectivity is associated with higher internalizing symptoms supports the idea that certain neural patterns, including those involved in spontaneous and off-task thought, may be linked to emotional difficulties (Kucyi et al., 2024). This finding is in line with previous work demonstrating that greater functional connectivity among key nodes within these networks at age seven was associated with depression and anxiety symptoms four years later (Whitfield-Gabrieli et al., 2020). Interestingly, however, we did not replicate this association longitudinally among the children above poverty. It is possible that prior work may have recruited a population with generally higher internalizing symptoms, or that we would see the same effects in our study emerge over a longer time delay. Cross-sectionally, our findings are also consistent with prior research suggesting that dysregulated connectivity within the DMN is implicated in anxiety and depression (Greicius et al., 2003; Sheline et al., 2010). Our study adds to this body of literature by showing that this relationship is particularly predictive of future internalizing symptoms for children living in poverty, emphasizing the potential role of socioeconomic factors in neural development and emotional health (Pollak & Wolfe, 2020).

However, our previous work with the same sample also found that higher LFPN-DMN connectivity was associated with better academic performance in children living in poverty. This finding might seem contradictory given the association of higher LFPN-DMN connectivity with higher internalizing symptoms. A possible explanation is that for children in poverty, the integration of these networks supports the cognitive resources necessary for academic success, but also puts them at risk for later internalizing symptoms. Previous research has indicated that lower LFPN-DMN connectivity, which reflects clearer segregation of task-positive and task-negative networks, is typically associated with better focus and cognitive performance, (Kelly et al., 2008; Hampson et al., 2010; Barber et al., 2013; Keller et al., 2015). However, in children from lower-income backgrounds, the opposite pattern was observed: higher LFPN-DMN connectivity was linked to better cognitive performance, possibly because it reflects an adaptive mechanism that helps these children cope with the high demands and stressors of their environments (Ellwood-Lowe et al., 2021). This heightened network integration might be beneficial for managing schoolwork in challenging circumstances, but it could also increase sensitivity to stress and emotional challenges, leading to higher internalizing symptoms.

The findings from this study provide critical insights into the intricate interplay between academic performance, brain connectivity, and mental health, particularly in the context of socioeconomic adversity. Our results suggest that for children living in poverty, academic performance is more closely tied to mental health outcomes, with poor grades being more strongly linked to internalizing symptoms such as anxiety and depression. This underscores the need for targeted interventions that address both educational and mental health support, especially for socioeconomically disadvantaged children. Our findings reveal that while higher grades do not predict future internalizing symptoms, the neural patterns linked to academic success, specifically higher LFPN-DMN connectivity, may predict such symptoms in children living in poverty. This underscores the complexity of interpreting neural connectivity as a marker for future mental health, particularly in different socioeconomic contexts. The association between higher LFPN-DMN connectivity and higher internalizing symptoms highlights a potential neural mechanism underlying these emotional difficulties. Importantly, this connectivity not only correlates with current internalizing symptoms but also predicts future symptoms for children living in poverty. This finding points to the importance of early detection and intervention strategies that incorporate neural markers, particularly as our results suggest that LFPN-DMN connectivity adds significant predictive value for future internalizing symptoms beyond what can be predicted from current symptoms alone. This aligns with the growing evidence that neural data can provide critical predictive insights into various domains, including education, performance, and health outcomes (Gabrieli et al., 2015).

Moreover, our study adds to the literature highlighting the significant impact of poverty on children’s emotional and cognitive development (Johnson et al., 2016; Farah, 2017; Lipina & Evers, 2017; Merz et al., 2020; Pollak & Wolfe, 2020; Noble et al., 2021). Socioeconomic status affects how academic performance relates to mental health, suggesting that children in poverty may experience a unique and more severe set of challenges. This calls for comprehensive policies that integrate educational, social, and healthcare resources to support the holistic development of children from low-income backgrounds.

## Limitations and Future Directions

Although this study provided valuable insights into the relationship between academic achievement, brain function, and mental health symptoms in children living in poverty, there are some limitations that should be addressed in future research. One of the limitations is that the directionality of the association between grades and internalizing symptoms is unclear. While the current findings suggest that worse grades may be a risk factor for internalizing symptoms, particularly in children living below poverty, it is also possible that internalizing symptoms could be a risk factor for worse grades. Indeed, one longitudinal study in Ireland following students in poverty found that 9 year-olds’ socioemotional problems negatively predicted their future math performance, along with several school and parent-related factors (Sheehan & Hadfield, 2024). Further exploration of the directionality of this association in the United States could be addressed in a prospective longitudinal study.

We were fortunate to have access to the ABCD study dataset, which allowed us to examine our questions in a large sample of children living in poverty. However, due to data anonymity concerns, poverty measures were estimated rather than directly measured. While these estimates can provide useful information about socioeconomic status, they may not fully capture the complexity of poverty and its effects on children’s lives. Future research could benefit from using more comprehensive measures of SES, such as those that take into account family income, education, and employment status. Poverty is also only a proxy for a vast number of possible experiences; it would be beneficial to explore these experiences directly in future work.

Finally, as mentioned above, it would be interesting to investigate the role of other brain networks in the relationship between academic performance, neural connectivity, and mental health outcomes. In addition, the boundaries between specific networks vary systematically across individuals, and it will be important to determine whether these also vary as a function of poverty status, perhaps shedding more insight into the results reported here. Finally, examining how other contextual factors, such as parenting styles, school environment, and community resources, moderate or mediate this relationship could provide further insights into how academic achievement, brain function, and mental health are interrelated in children living in poverty.

## Conclusions

Overall, this set of findings underscore the significance of going beyond convenience samples to better understand the diverse experiences of children in poverty. They highlight the critical need for educational systems to recognize and address the dual burden of academic and emotional challenges faced by children in poverty. By fostering environments that support both academic success and emotional well-being, we can help mitigate the long-term adverse effects of poverty on children’s lives. Investing in such integrative approaches not only benefits individual children but also promotes greater societal equity and well-being. This research contributes to a deeper understanding of the socioeconomic determinants of mental health and academic success, advocating for a multidimensional approach to child development that can inform future educational and public health policies.

## Acknowledgments

This study relied on data from the ABCD study (see below). The extensive contributions of the large team of ABCD leaders and organizers, staff and data curators, as well as the participation of families and children, were essential in making this study possible. SP was supported by UC Berkeley’s Regents’ and Chancellors’ Scholarship, Robert and Colleen Haas Scholars Program Fellowship, Biology Scholars Program Research Fellowship, and Summer Undergraduate Research Fellowship. Data used in the preparation of this article were obtained from the Adolescent Brain Cognitive Development^SM^ (ABCD) Study (https://abcdstudy.org), held in the NIMH Data Archive (NDA). This is a multisite, longitudinal study designed to recruit more than 10,000 children age 9-10 and follow them over 10 years into early adulthood. The ABCD Study® is supported by the National Institutes of Health and additional federal partners under award numbers U01DA041048, U01DA050989, U01DA051016, U01DA041022, U01DA051018, U01DA051037, U01DA050987, U01DA041174, U01DA041106, U01DA041117, U01DA041028, U01DA041134, U01DA050988, U01DA051039, U01DA041156, U01DA041025, U01DA041120, U01DA051038, U01DA041148, U01DA041093, U01DA041089, U24DA041123, U24DA041147. A full list of supporters is available at https://abcdstudy.org/federal-partners.html. A listing of participating sites and a complete listing of the study investigators can be found at https://abcdstudy.org/consortium_members/. ABCD consortium investigators designed and implemented the study and/or provided data but did not necessarily participate in the analysis or writing of this report. This manuscript reflects the views of the authors and may not reflect the opinions or views of the NIH or ABCD consortium investigators. The ABCD data repository grows and changes over time. The ABCD data used in this report came from 10.15154/8873-zj65.

