## Supplemental Material for "Academic Success and Mental Health: The Paradox of Frontoparietal-Default Mode Network Coupling among Children Facing Poverty"

### Table of Contents


| | |
|---|---|
| **Cross-Sectional Analyses: Associations with Internalizing Subscales** | **2-10** |
| Children's Grades and Internalizing Subscales | **2-4** |
| Children's LFPN-DMN Connectivity Levels and Internalizing Subscales | **5-7** |
| Children's CON-DMN Connectivity Levels and Internalizing Subscales | **7-9** |
| Children's CON-LFPN Connectivity Levels and Internalizing Subscales | **9-10** |
| **Longitudinal Analyses: Associations Over Time** | **11-22** |
| Children's Grades and Internalizing Symptoms Over Time | **11-14** |
| Children's LFPN-DMN Connectivity Levels and Internalizing Symptoms Over Time | **15-17** |
| Children's CON-DMN Connectivity Levels and Internalizing Symptoms Over Time | **17-20** |
| Children's CON-LFPN Connectivity Levels and Internalizing Symptoms Over Time | **20-22** |

### Cross-Sectional Analyses

#### *Association Between Children's Grades and Internalizing Subscales*

*Anxiety/Depression Symptoms***.** As shown in Figure S1A, mirroring results for the composite internalizing scores, worse grades were associated with greater anxiety/depression symptoms ($B$ = 2.51, SE = 0.86; $X^2$ (4) = 163.59, $p$ < .001). The association between anxiety/depression symptoms and grades varied as a function of whether or not children were in poverty ($X^2$ (4) = 15.03, $p$ = .005). Specifically, while children both above and below poverty with worse grades typically had higher anxiety/depression symptoms, this association was more positive for children below poverty ($B$ = 6.02, SE = 0.86 ; $X^2$ (4) = 59.77, $p$ < .001) than for children above poverty ($B$ = 2.55, SE = 0.95; $X^2$ (4) = 120.73, $p$ < .001), mirroring results with the internalizing composite.

*Withdrawal/Depression Symptoms.* As shown in Figure S1B, across the group, worse grades were similarly associated with greater withdrawal/depression symptoms ($B$ = 2.61, SE = 0.83; $X^2$ (4) = 221.60, $p$ < .001). Children in poverty experienced greater withdrawal/depression symptoms on average than children above poverty ($B$ = 2.53, SE = 0.40; $X^2$ (1) = 58.74, $p$ < .001). In addition, the association between withdrawal/depression symptoms and grades varied as a function of whether children were in poverty ($X^2$ (4) = 13.72, $p$ = .008). Specifically, while children both above and below poverty with worse grades typically had higher withdrawal/depression symptoms, this association between grades and internalizing symptoms was more positive for children below poverty ($B$ = 6.64, SE = 1.01; $X^2$ (4) = 75.56, $p$ < .001) than for children above poverty ($B$ = 2.66, SE = 0.77; $X^2$ (4) = 133.40, $p$ < .001), mirroring results above.

*Somatic Symptoms*. As shown in Figure S1C, worse grades were similarly associated with higher somatic symptoms (B = 0.53, SE = 0.87; $X^2$ (4) = 53.62, p < .001). Children in poverty experienced greater somatic symptoms on average than children above poverty (B = 1.00, SE = 0.43; $X^2$ (1) = 8.72, p = .003). However, in contrast to the prior results, the interaction did not reach significance ($X^2$ (4) = 4.00; p = 0.406), indicating that the association was consistent regardless of poverty status.

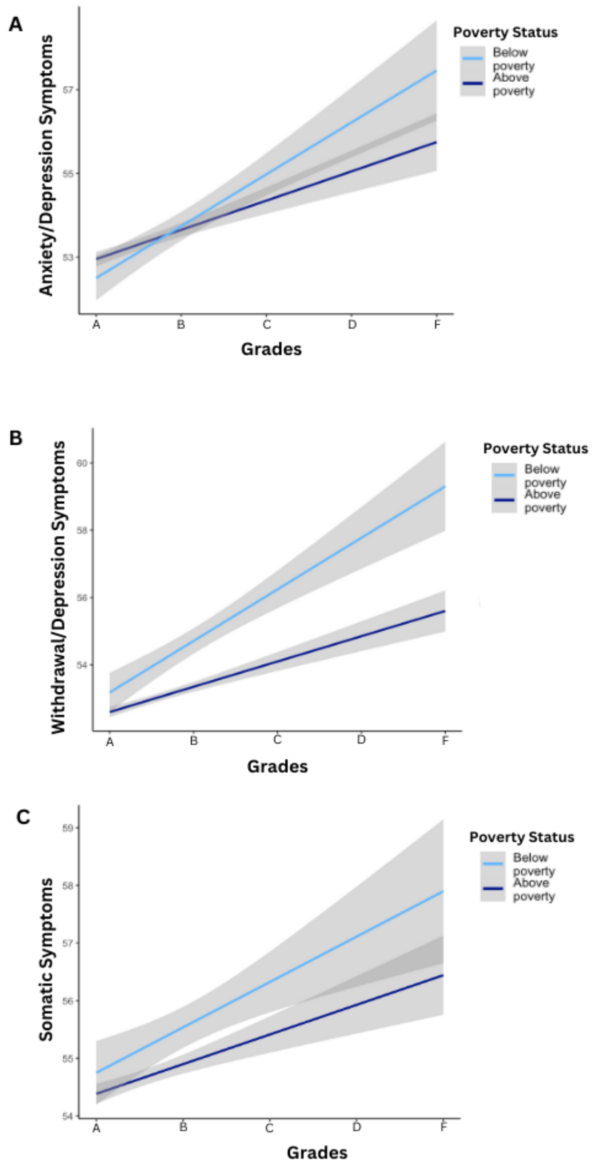

**Fig. S1 Association Between Children's Grades in School and Internalizing Subscales.** Relation between children's school grades and their (A) anxiety/depression symptoms, (B) withdrawal/depression symptoms, and (C) somatic symptoms for children below poverty (light blue) and children above poverty (dark blue). The gray bars indicate mean values with ±95% confidence intervals for a linear model.

***Association Between Children's LFPN-DMN Connectivity Levels and Internalizing Subscales***

*Anxiety/Depression Symptoms.* As shown in Figure S2A, there was a main effect of LFPN-DMN connectivity ($B$ = 4.99, SE = 1.61; $X^2$ (1) = 9.01, $p$ = .003). However, the interaction effect between poverty status and LFPN-DMN connectivity in relation to anxiety/depression symptoms was not statistically significant ($X^2$ (1) = 0.81 , $p$ = .369), indicating that the association was consistent regardless of poverty status.

*Withdrawal/Depression Symptoms.* As shown in Figure S2B, there was a main effect of LFPN-DMN connectivity ($B$ = 5.21, SE = 1.52; $X^2$ (1) = 17.73, $p$ < .001). However, the interaction effect between poverty status and LFPN-DMN connectivity in relation to withdrawal/depression symptoms was not statistically significant ($X^2$ (1) = 1.04 , $p$ = 0.309), indicating that the association between LFPN-DMN connectivity and withdrawal/depression symptoms was consistent regardless of poverty status.

*Somatic Symptoms.* As shown in Figure S2C, the main effect of LFPN-DMN connectivity was non-significant ($B$ = 2.28, SE = 1.61; $X^2$ (1) = 0.75, $p$ = .388) nor was the interaction significant ($X^2$ (1) = 2.37, $p$ = 0.124).

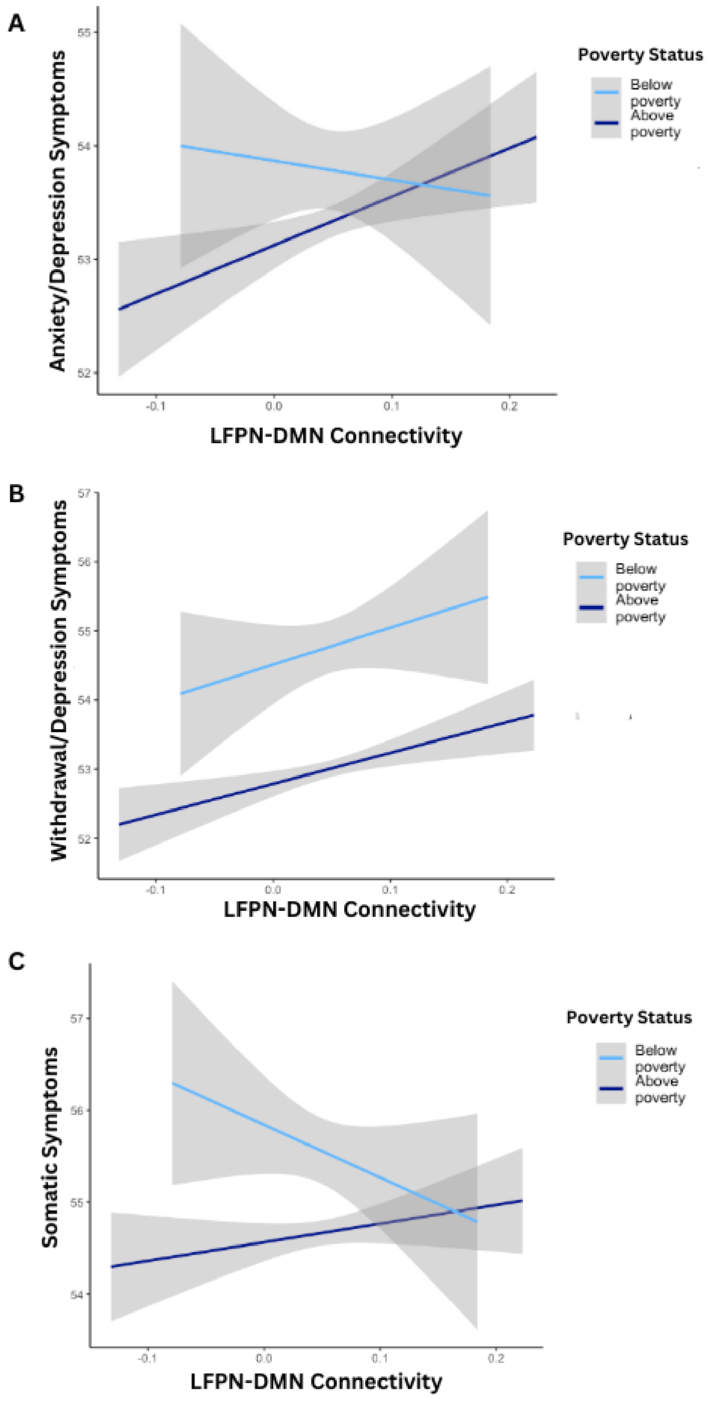


**Fig. S2 Association Between Children's LFPN-DMN Connectivity Levels and Internalizing Subscales.** Relation between children's LFPN-DMN connectivity and their (A) anxiety/depression symptoms, (B) withdrawal/depression symptoms, and (C) somatic symptoms for children below poverty (light blue) and children above poverty (dark blue). The gray bars indicate mean values with ±95% confidence intervals for a linear model.

#### *Association Between Children's CON-DMN Connectivity Levels and Internalizing Subscales*

*Anxiety/Depression Symptoms.* Children's CON-DMN connectivity levels exhibited a significant association with their anxiety/depression symptoms ($B = 1.86$, SE = 1.30; $X^2(1) = 4.17$, $p = .041$). Importantly, there was no significant interaction between CON-DMN connectivity and poverty status ($X^2(1) = 1.36$, $p = .243$), indicating consistent associations across these subgroups.

*Withdrawal/Depression Symptoms.* There was a significant association between children's CON-DMN connectivity levels and their withdrawal/depression symptoms ($B = 4.42$, SE = 1.22; $X^2(1) = 20.64$, $p < .001$), indicating that higher CON-DMN connectivity was linked to more significant withdrawal/depression symptoms. The interaction analysis did not reach statistical significance ($X^2(1) = 2.27$, $p = .132$), suggesting that the association between CON-DMN connectivity levels and withdrawal/depression symptoms did not significantly differ between children below and above the poverty line.

*Somatic Symptoms.* CON-DMN connectivity levels did not show a significant association with their somatic symptoms ($B = -1.09$, SE = 1.30, $X^2(1) = 0.03$, $p = .858$) nor was the interaction significant ($X^2(1) = 3.05$, $p = .081$).

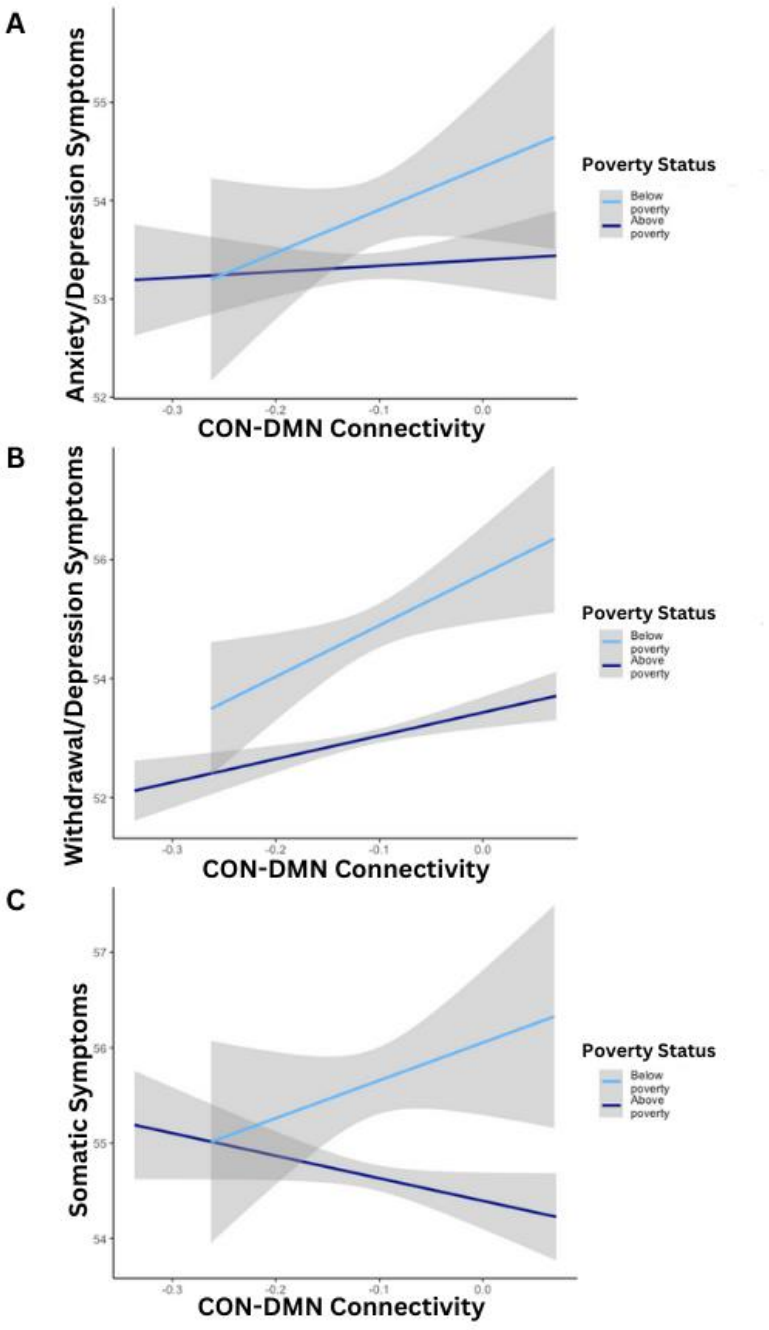


**Fig. S3 Association Between Children's CON-DMN Connectivity Levels and Internalizing Subscales.** Relation between children's CON-DMN connectivity and their (A) anxiety/depression symptoms, (B) withdrawal/depression symptoms, and (C) somatic symptoms for children below poverty (light blue) and children above poverty (dark blue). The gray bars indicate mean values with ±95% confidence intervals for a linear model.

***Association Between Children's CON-LFPN Connectivity Levels and Internalizing Subscales***

*Anxiety/Depression Symptoms.* Children's CON-LFPN connectivity levels showed a non-significant association with their anxiety/depression symptoms (*B* = -0.12, SE = 1.53; $X^2$ (1) = 0.21, *p* = .645), indicating no substantial effect. Importantly, the interaction analysis also did not reach statistical significance ($X^2$ (1) = 1.67, *p* = .196), suggesting that the association between CON-LFPN connectivity levels and anxiety/depression symptoms was consistent across both poverty and non-poverty populations.

*Withdrawal/Depression Symptoms.* Children's CON-LFPN connectivity levels showed no significant association with their withdrawal/depression symptoms (*B* = -1.16, SE = 1.44; $X^2$ (1) = 0.07, *p* = .784)Importantly, the interaction analysis reached statistical significance ($X^2$ (1) = 7.34, *p* = 0.007), indicating that the association between CON-LFPN connectivity levels and withdrawal/depression symptoms significantly differed between children below and above the poverty line. Breaking down this interaction revealed that among children below the poverty line higher CON-LFPN connectivity was related to less pronounced withdrawal/depression symptoms (*B* = -0.72, SE = 1.34; $X^2$ (1) = 0.29, *p* = .591) compared to children above poverty (*B* = 8.25, SE = 4.27; $X^2$ (1) = 3.74, *p* = .053), though these directional associations were non-significant.

*Somatic Symptoms.* Children's CON-LFPN connectivity levels displayed no significant association with their somatic symptoms (*B* = 0.22, SE = 1.53; $X^2$ (1) = 0.24, *p* = .625). Importantly, the interaction analysis did not reach statistical significance ($X^2$ (1) = 0.61, p = .435), indicating that the association between CON-LFPN connectivity levels and somatic symptoms was consistent across both poverty and non-poverty populations.

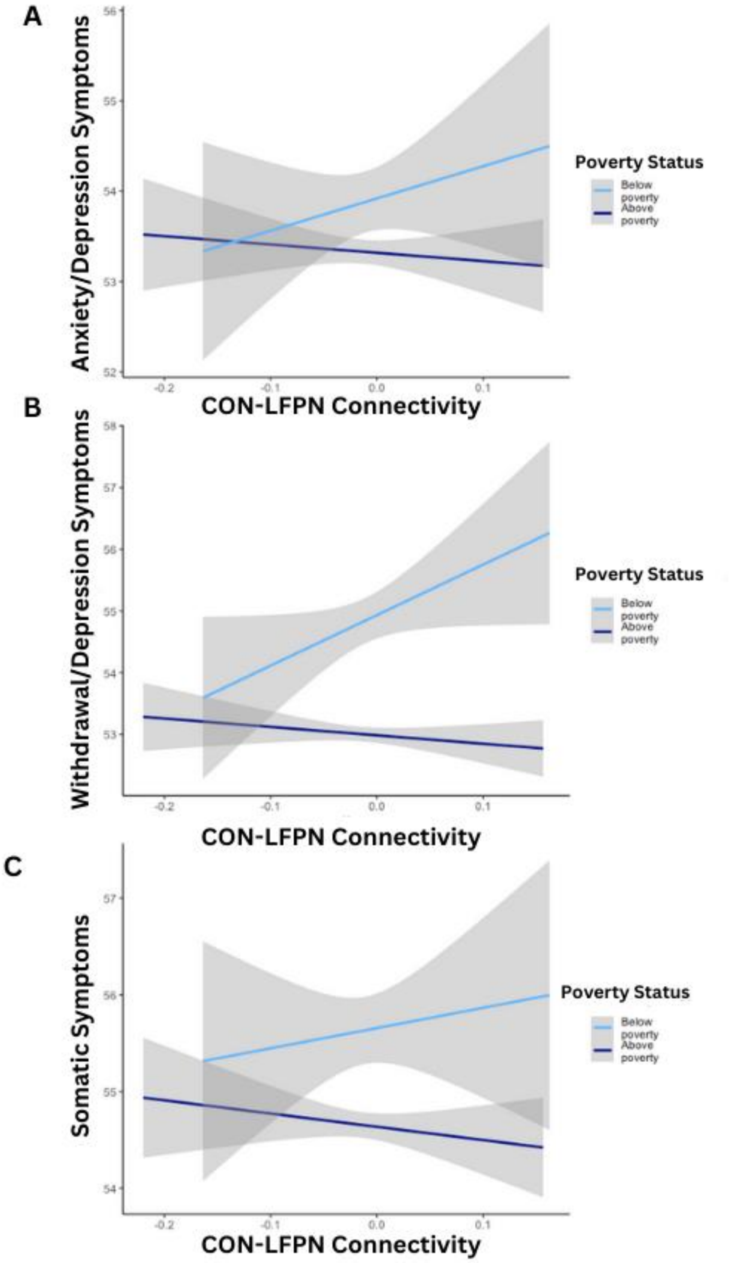


**Fig. S4 Association Between Children's CON-LFPN Connectivity Levels and Internalizing Subscales.** Relation between children's CON-LFPN connectivity and their (A) anxiety/depression symptoms, (B) withdrawal/depression symptoms, and (C) somatic symptoms for children below poverty (light blue) and children above poverty (dark blue). The gray bars indicate mean values with ±95% confidence intervals for a linear model.

### Longitudinal Analyses

To understand the evolution of the interaction between poverty status and internalizing symptoms in predicting grades, LFPN-DMN, CON-DMN, and CON-LFPN connectivity we performed linear mixed-effects models, using the lme4 package to separately test the relation between (1) grades, (2) LFPN-DMN, (3) CON-DMN, and (4) CON-LFPN connectivity and internalizing symptoms, including a time point interaction with grades or connectivity (as appropriate), across four time points: (1) baseline, (2) 1-year follow-up, (3) 2-year follow-up, and (4) 3-year follow-up. All models included an interaction of poverty level, fixed effects for child age, and random intercepts for site and family; LFPN-DMN, CON-DMN, and CON-LFPN models additionally included fixed effects for head motion (mean framewise displacement).

#### *Association Between Children's Grades and Internalizing Symptoms Over Time*

Across the entire group and all timepoints, lower grades were associated with greater internalizing symptoms ($B$ = 2.11, SE = 0.58, $X^2$ (1) = 337.20, $p$ < .001). The main effect of time was also significant across all timepoints ($X^2$ (3) = 17.19, $p$ < .001) indicating that children's level of internalizing symptoms changed over early adolescence. In addition, the three-way interaction of grades, poverty, and time was significant ($X^2$ (12) = 25.29, $p$ = .013) indicating that the impact of grades on internalizing over time differs for children above and below the poverty line. To gain a deeper understanding of the relationship between children's grades and internalizing symptoms as it evolves over time, we analyzed each time point separately. Contrary to hypotheses, the link between grades and internalizing symptoms was stronger among children below the poverty line across all timepoints.

**Table S1A. Baseline**

| | B | SE | $X^2$ | p |
|---|---|---|---|---|
| **Grades** | 2.00 | 0.78 | 195.55 | < .001 *** |
| **Poverty Status** | 1.55 | 0.38 | 8.79 | .003 ** |
| **Grades:Poverty Status** | 3.69 | 1.08 | 14.18 | .007 ** |
| **Above Poverty** | 2.05 | 0.75 | 137.29 | < .001 *** |
| **Below Poverty** | 5.94 | 0.87 | 64.72 | < .001 *** |

**Table S1A.** Results of linear mixed effects model associating internalizing symptoms with a two-way interaction between grades at baseline and poverty status. Chi-squared and significance values from Type II anova, using Anova function in car.

**Table S1B. One-Year Follow-Up**

| | B | SE | $X^2$ | p |
|---|---|---|---|---|
| **Grades** | 3.80 | 0.83 | 173.25 | < .001 *** |
| **Poverty Status** | 0.60 | 0.39 | 0.85 | .355 |
| **Grades:Poverty Status** | 1.36 | 1.10 | 3.38 | .497 |

**Table S1B.** Results of linear mixed effects model associating internalizing symptoms with a two-way interaction between grades at T1 and poverty status. Chi-squared and significance values from Type II anova, using Anova function in car.

**Table S1C. Two-Year Follow-Up**

| | B | SE | $X^2$ | p |
|---|---|---|---|---|
| **Grades** | 3.01 | 0.80 | 114.70 | < .001 *** |
| **Poverty Status** | 1.18 | 0.44 | 0.005 | .946 |
| **Grades:Poverty Status** | 3.26 | 1.24 | 13.97 | .007 ** |
| **Above Poverty** | 3.00 | 0.78 | 69.43 | < .001 *** |
| **Below Poverty** | 6.26 | 1.07 | 48.39 | < .001 *** |

**Table S1C**. Results of linear mixed effects model associating internalizing symptoms with a two-way interaction between grades at T2 and poverty status. Chi-squared and significance values from Type II anova, using Anova function in car.

**Table S1D. Three-Year Follow-Up**

| | B | SE | $X^2$ | p |
|---|---|---|---|---|
| **Grades** | 3.62 | 0.80 | 150.01 | < .001 *** |
| **Poverty Status** | 0.50 | 0.41 | 3.340 | .068 |
| **Grades:Poverty Status** | 2.62 | 1.13 | 12.08 | .017 * |
| **Above Poverty** | 3.59 | 0.75 | 92.22 | < .001 *** |
| **Below Poverty** | 6.22 | 0.92 | 60.56 | < .001 *** |

**Table S1D.** Results of linear mixed effects model associating internalizing symptoms with a two-way interaction between grades at T3 and poverty status. Chi-squared and significance values from Type II anova, using Anova function in *car*.

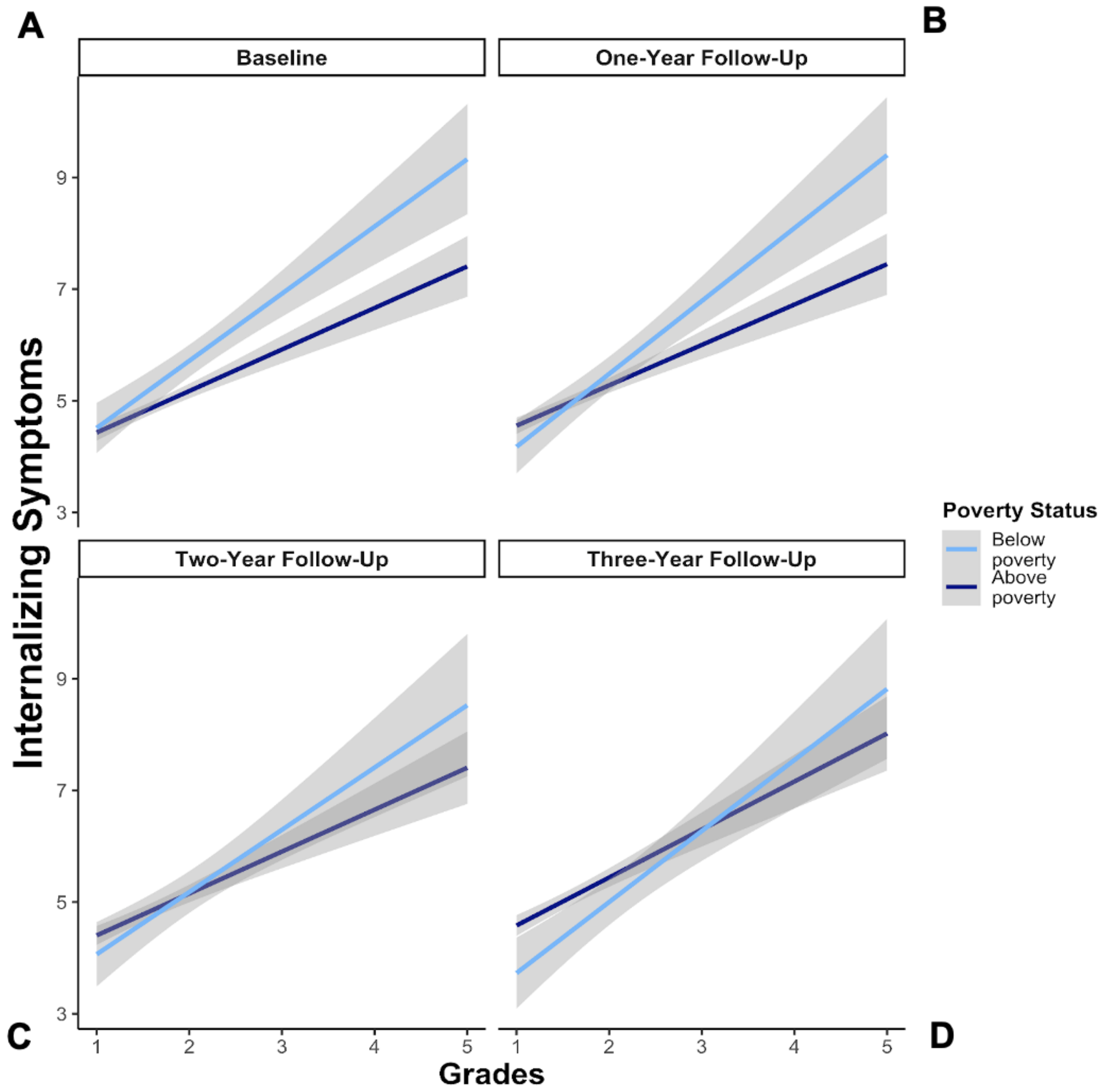


**Fig. S5 Association Between Children's Grades and Internalizing Symptoms Over Time.** Relations between children's grades and their internalizing symptoms at (A) baseline, (B) one-year follow-up, (C) two-year follow-up, and (D) three-year follow-up. Children below poverty (light blue) and children above poverty (dark blue). The gray bars indicate mean values with ±95% confidence intervals for a linear model.

***Association Between Children's LFPN-DMN Connectivity Levels and Internalizing Symptoms Over Time***

Higher LFPN-DMN connectivity was positively associated with more internalizing symptoms ($B$ = 3.94, SE = 1.30, $X^2$ (1) = 4.46, $p$ = 0.035), supporting our hypotheses. There was also a significant interaction between LFPN-DMN connectivity and time ($X^2$ (1) = 4.38, $p$ = .036); follow-up analyses revealed that the concurrent association between LFPN-DMN connectivity and internalizing symptoms was significant at baseline but not at the two-year follow-up (Table S2B). Thus, children's LFPN-DMN connectivity at T2 was not related to their internalizing symptoms at the same time point.

The interaction between LFPN-DMN and time was significant ($X^2$ (1) = 4.38, $p$ = .036). Additionally, the interaction between poverty and time was significant ($X^2$ (1) = 10.80, $p$ = .001) suggesting that the influence of poverty on children's internalizing symptoms varies over time. Importantly, the three way interaction of LFPN-DMN connectivity, poverty, and time was not significant ($X^2$ (1) = 0.12, $p$ = .728). To gain a deeper understanding of these relationships, we analyzed each time point separately.

Results at baseline revealed a significant positive association, supporting the hypothesis that higher LFPN-DMN connectivity corresponds to greater internalizing symptoms. However, the interaction with poverty status did not reach significance suggesting a consistent association across economic backgrounds. However, at the two-year follow-up, the significant association between LFPN-DMN connectivity and internalizing symptoms observed at baseline diminished.

**Table S2A. Baseline**

| | B | SE | $X^2$ | p |
|---|---|---|---|---|
| **LFPN-DMN Connectivity** | 4.61 | 1.45 | 9.88 | .002 ** |
| **Poverty Status** | 1.22 | 0.24 | 42.48 | < .003 *** |
| **LFPN-DMN Connectivity:Poverty Status** | -2.66 | 3.54 | 0.57 | .451 |

**Table S2A.** Results of linear mixed effects model associating internalizing symptoms with a two-way interaction between LFPN-DMN connectivity at baseline and poverty status. Chi-squared and significance values from Type II anova, using Anova function in car.

**Table S2B. Two-Year Follow-Up**

| | B | SE | $X^2$ | p |
|---|---|---|---|---|
| **LFPN-DMN Connectivity** | 1.48 | 1.84 | 0.42 | .516 |
| **Poverty Status** | 0.35 | 0.32 | 1.08 | .300 |
| **LFPN-DMN Connectivity:Poverty Status** | -2.53 | 4.70 | 0.29 | .590 |

**Table S2B.** Results of linear mixed effects model associating internalizing symptoms with a two-way interaction between LFPN-DMN connectivity at T2 and poverty status. Chi-squared and significance values from Type II anova, using Anova function in car.

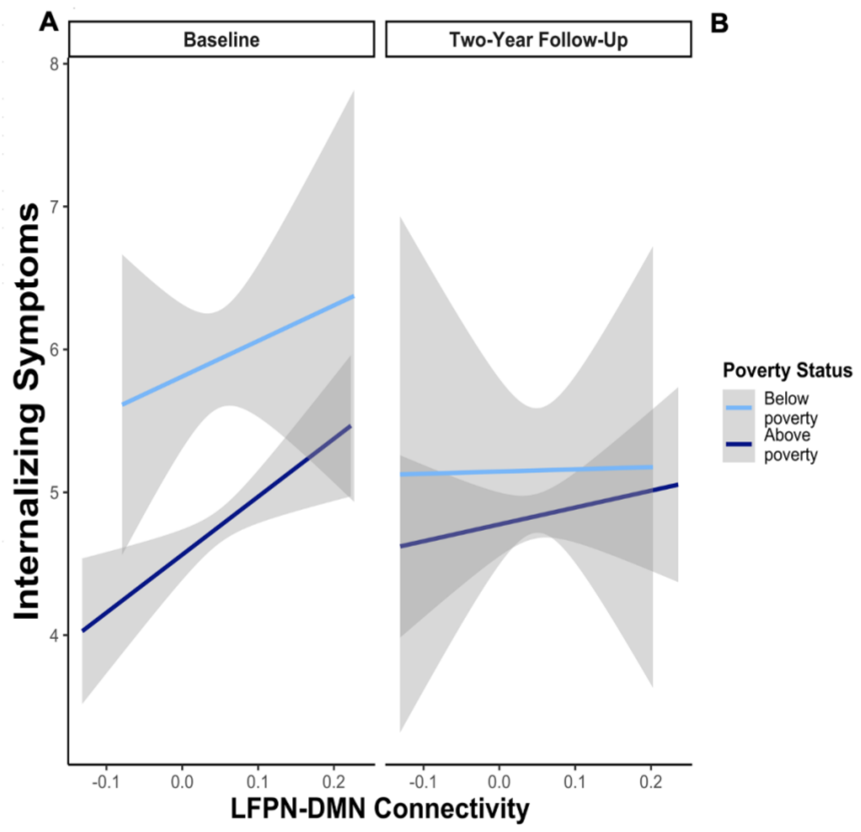


**Fig. S6 Association Between Children's LFPN-DMN Connectivity and Internalizing Symptoms Over Time.** Relations between children's LFPN-DMN connectivity and their internalizing symptoms at (A) baseline, and (B) two-year follow-up. Children below poverty (light blue) and children above poverty (dark blue). The gray bars indicate mean values with ±95% confidence intervals for a linear model.

#### *Association Between Children's CON-DMN Connectivity Levels and Internalizing Symptoms Over Time*

Across both time points (T0 and T2), we observed that children's CON-DMN connectivity levels did not exhibit a significant association with their internalizing symptoms (*B* = 0.57, SE = 1.05, $X^2$ (1) = 0.05, *p* = 0.82). Notably, the main effect of time was found to be significant across all timepoints ($X^2$ (1) = 9.92, *p* = .002), indicating a consistent shift in internalizing symptoms over time. The only significant interaction we found was between poverty and time ($X^2$ (1) = 10.24, $p$ = .001). To gain a deeper understanding of these relationships, we analyzed each time point separately.

At baseline, no significant association was found between CON-DMN connectivity and internalizing symptoms. Notably, poverty status independently influenced internalizing symptoms, with children in poverty reporting higher levels than their counterparts. The interaction between CON-DMN connectivity and poverty status did not reach significance. Similarly, at the two-year follow-Up, CON-DMN connectivity showed no significant association with internalizing symptoms, and poverty status continued to be a significant factor. The interaction between CON-DMN connectivity and poverty status remained non-significant. The findings suggest a consistent lack of direct association between CON-DMN connectivity and internalizing symptoms, highlighting the robust impact of poverty status on children's mental health outcomes.

**Table S3A. Baseline**

| | **B** | **SE** | **$X^2$** | **p** |
|---|---|---|---|---|
| **CON-DMN Connectivity** | 0.98 | 1.17 | 2.60 | .107 |
| **Poverty Status** | 1.60 | 0.34 | 42.13 | < .001 *** |
| **CON-DMN Connectivity:Poverty Status** | 4.78 | 2.80 | 2.91 | .088 |

**Table S3A.** Results of linear mixed effects model associating internalizing symptoms with a two-way interaction between CON-DMN connectivity at baseline and poverty status. Chi-squared and significance values from Type II anova, using Anova function in car.

**Table S3B. Two-Year Follow-Up**

| | **B** | **SE** | **$X^2$** | **p** |
|---|---|---|---|---|
| **CON-DMN Connectivity** | -1.63 | 1.48 | 2.25 | .134 |

| | | | | |
|---|---|---|---|---|
| **Poverty Status** | -0.12 | 0.47 | 1.15 | .284 |
| **CON-DMN Connectivity:Poverty Status** | -3.17 | 3.76 | 0.71 | .399 |

**Table S3B.** Results of linear mixed effects model associating internalizing symptoms with a two-way interaction between CON-DMN connectivity at T2 and poverty status. Chi-squared and significance values from Type II anova, using Anova function in car.

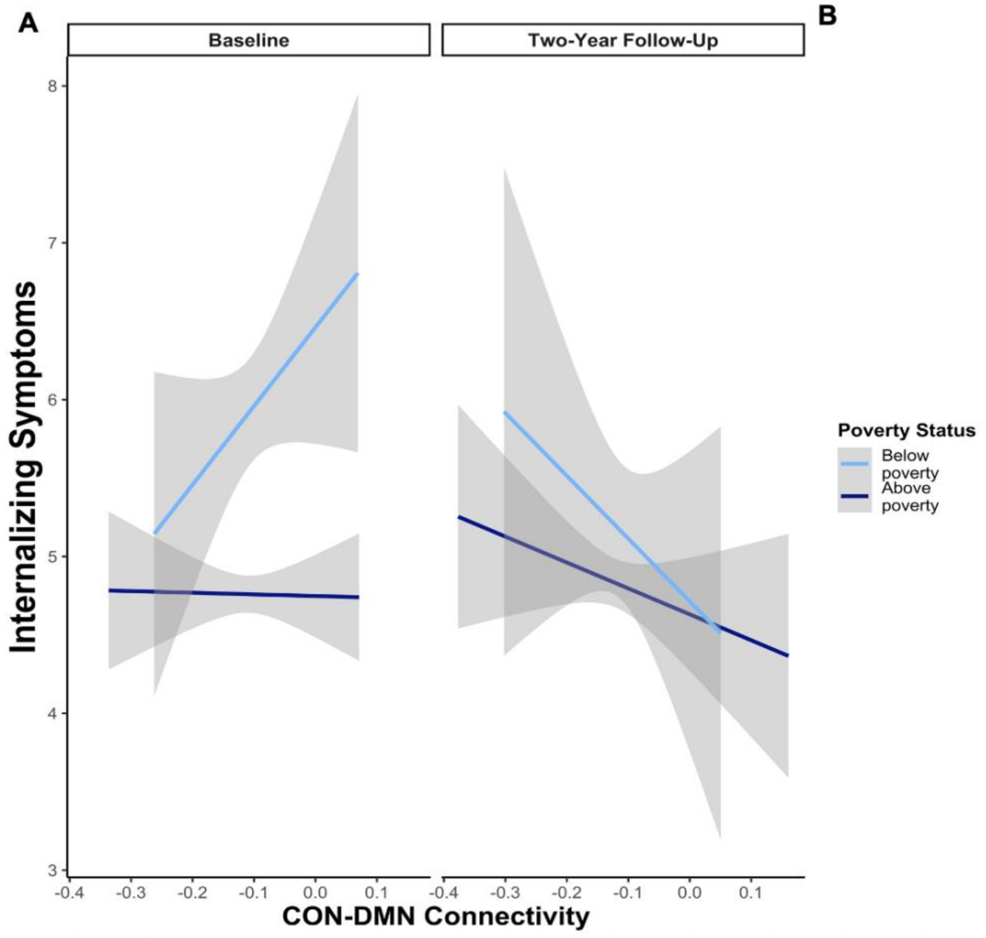


**Fig. S7 Association Between Children's CON-DMN Connectivity and Internalizing Symptoms Over Time.** Relations between children's CON-DMN connectivity and their internalizing symptoms at (A) baseline, and (B) two-year follow-up. Children below poverty (light blue) and children above poverty (dark blue). The gray bars indicate mean values with ±95% confidence intervals for a linear model.

***Association Between Children's CON-LFPN Connectivity Levels and Internalizing Symptoms Over Time***

Across all time points, we observed that children's CON-LFPN connectivity levels did not exhibit a significant association with their internalizing symptoms ($B$ = -0.50, SE = 1.24, $X^2$ (1) = 0.18, $p$ = 0.67). Notably, the main effect of time was found to be significant across all timepoints ($X^2$ (1) = 9.92, $p$ = .002), indicating a consistent shift in internalizing symptoms over time. The only significant interaction we found was between poverty and time ($X^2$ (1) = 10.81, $p$ = .001). To gain a deeper understanding of these relationships, we analyzed each time point separately.

At baseline, there was no significant association between CON-LFPN connectivity and internalizing symptoms. A trend in the interaction between CON-LFPN connectivity and poverty status hinted at a potential interplay, though it did not reach statistical significance. Similarly, at the two-year follow-up, no significant association was found between CON-LFPN connectivity levels and internalizing symptoms. The interaction between CON-LFPN connectivity and children's poverty status also did not reach significance, suggesting a consistent association regardless of economic background, contrary to initial hypotheses.

**Table S4A. Baseline**

| | **B** | **SE** | **$X^2$** | **p** |
|---|---|---|---|---|
| **CON-LFPN Connectivity** | -0.43 | 1.38 | 0.23 | .628 |
| **Poverty Status** | 1.16 | 0.17 | 41.91 | < .001 *** |
| **CON-LFPN Connectivity:Poverty Status** | 6.69 | 3.42 | 3.81 | .051 |

**Table 4A.** Results of linear mixed effects model associating internalizing symptoms with a two-way interaction between CON-LFPN connectivity at baseline and poverty status. Chi-squared and significance values from Type II anova, using Anova function in car.

**Table S4B. Two-Year Follow-Up**

| | B | SE | $X^2$ | p |
|---|---|---|---|---|
| **CON-LFPN Connectivity** | -0.54 | 1.76 | 0.07 | .798 |
| **Poverty Status** | 0.24 | 0.23 | 1.09 | .297 |
| **CON-LFPN Connectivity:Poverty Status** | 0.94 | 4.78 | 0.04 | .844 |

**Table 4B.** Results of linear mixed effects model associating internalizing symptoms with a two-way interaction between CON-LFPN connectivity at T2 and poverty status. Chi-squared and significance values from Type II anova, using Anova function in car.

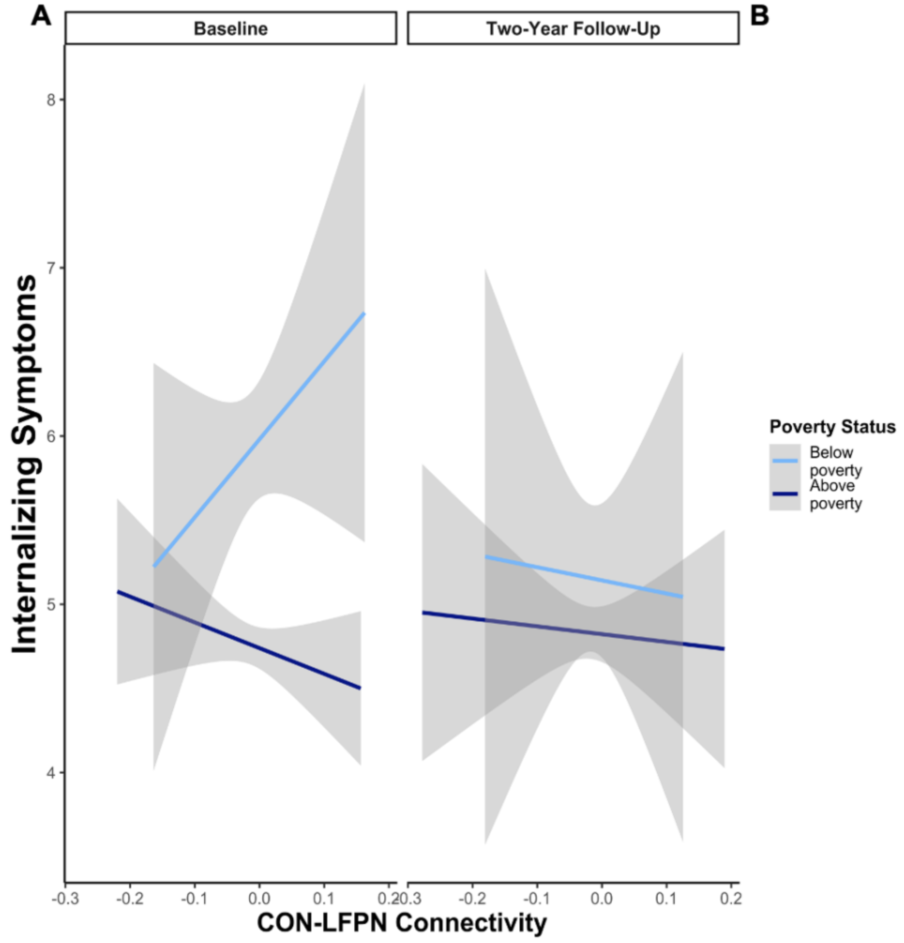


**Fig. S8 Association Between Children's CON-LFPN Connectivity and Internalizing Symptoms Over Time.** Relations between children's CON-LFPN connectivity and their internalizing symptoms at (A) baseline, and (B) two-year follow-up. Children below poverty (light blue) and children above poverty (dark blue). The gray bars indicate mean values with ±95% confidence intervals for a linear model.
